# Common low complexity regions for SARS-CoV-2 and human proteomes as potential multidirectional risk factor in vaccine development

**DOI:** 10.1101/2020.08.11.245993

**Authors:** Aleksandra Gruca, Joanna Ziemska-Legiecka, Patryk Jarnot, Elzbieta Sarnowska, Tomasz J. Sarnowski, Marcin Grynberg

## Abstract

The rapid spread of the COVID-19 demands immediate response from the scientific communities. Appropriate countermeasures mean thoughtful and educated choice of viral targets (epitopes). There are several articles that discuss such choices in the SARS-CoV-2 proteome, other focus on phylogenetic traits and history of the Coronaviridae genome/proteome. However none consider viral protein low complexity regions (LCRs). Recently we created the first methods that are able to compare such fragments. We show that five low complexity regions (LCRs) in three proteins (nsp3, S and N) encoded by the SARS-CoV-2 genome are highly similar to regions from human proteome. As many as 21 predicted T-cell epitopes and 27 predicted B-cell epitopes overlap with the five SARS-CoV-2 LCRs similar to human proteins. Interestingly, replication proteins encoded in the central part of viral RNA are devoid of LCRs. Similarity of SARS-CoV-2 LCRs to human proteins may have implications on the ability of the virus to counteract immune defenses. The vaccine targeted LCRs may potentially be ineffective or alternatively lead to autoimmune diseases development. These findings are crucial to the process of selection of new epitopes for drugs or vaccines which should omit such regions.

**Author summary:** The outbreak of the COVID-19 disease affects humans all over the globe. More and more people get sick and many die because of the deadly SARS-CoV-2 virus. The whole machinery of this pathogen is enclosed in a short sequence of nucleotides, building blocks for both RNA and DNA strands. This RNA virus encodes less than 30 protein sequences that change the fate of our societies. Its proteins are composed of 20 amino acids (building bricks) that are usually used quite freely by proteins. However, there are fragments where only one or a few amino acids are used. We name those low complexity regions (LCRs). We invented the first programmes able to compare such LCRs. Using this new methodology we were able to show similarity of some viral proteins to human ones. This discovery has a serious implication when designing vaccines or drugs. It means that companies should not use these very LCRs as targets because it may trigger an autoimmune disease. On the other hand this specific similarity may suggest some kind of disguise of viral proteins into the machinery of human cells.

## Introduction

At the very end of 2019 the Chinese Center for Disease Control (China CDC) reported several severe pneumonia cases of unknown etiology in the city of Wuhan. The causative agent of the disease was a previously unknown *Betacoronavirus* named SARS-CoV-2. The virus quickly spread all over the globe (https://www.who.int/emergencies/diseases/novel-coronavirus-2019; https://www.worldometers.info/coronavirus/) and as of today (June 20202) the number of infections and the number of deaths were globally still on the rise.

Coronaviruses are widespread in vertebrates and cause a plethora of respiratory, enteric, hepatic, and neurologic issues. Some of the animal coronaviruses exhibited ability to transmit to human e.g. the severe acute respiratory syndrome coronavirus (SARS-CoV) in 2003 and Middle East respiratory syndrome coronavirus (MERS-CoV) in 2012 had caused human epidemics [1,2]. SARS-CoVs enters cells via the angiotensin-converting enzyme 2 (ACE2) receptor [3,4]. The SARS-CoV-2 first infects airways and binds to ACE2 on alveolar epithelial cells. Both viruses are potent inducers of inflammatory cytokines [5]. The virus activates immune cells and induces the secretion of inflammatory cytokines and chemokines into pulmonary vascular endothelial cells in a so-called “cytokine storm” [6–9]. Since most infected individuals are apparently asymptomatic it is hard to assess the prevalence of SARS-CoV-2 in global or even local populations. Lack of appropriate testing quantities also plays a role. To date there are no fully effective drugs or vaccines against SARS-CoV-2 [6,10–14].

Several recent articles have suggested that SARS-CoV-2 proteome fragments are important to the viral lifecycle, several of which are conserved in the Coronaviridae family and which are possible targets as epitopes [15–22]. This is of importance since the choice of attack points is crucial to the success of the therapy (drug design) or prevention (vaccine design).

The quest for optimal targets is difficult and focuses on both experimental and bioinformatic means. This may directly involve laboratory experiments or databases like the Immune Epitope Database and Analysis Resource (IEDB)[16,22,23]. The second approach uses the search for similarities with other viruses in order to identify conserved regions of the viral genome [19,24,25].

Although both methods focus on the viral RNA/proteome sequence, these approaches clearly treat this sequence as a whole. However, one can clearly distinguish between protein high complexity regions (HCRs) encompassing most of the genome and low complexity regions (LCRs). LCRs are often described as ‘unstructured’ or are simply not annotated. Our recent experiments however show that the search for similar sequences in LCRs is almost impossible using standard methods, like BLAST or HHblits [26,27]. This is why we created three algorithms that are able to compare low complexity protein sequences, GBSC, MotifLCR and LCR-BLAST [28–32].

The current situation due to COVID-19 is critical, and this work aims to compare the SARS-CoV-2 LCRs with the human proteome. In this work we show that numerous fragments of the viral proteome are very similar to the human ones. It has been recently shown that the furin cleavage site of Spike SARS-CoV-2 protein shares similarity with the human epithelial sodium channel [33]. Our findings suggest that identified fragments of spike, nsp3 and nucleocapsid proteins should not be considered as epitopes neither for vaccine nor drug design.

Moreover, our hypothesis is supported by the malaria molecular evolution and vaccine design study clearly indicating that LCRs may play a role in immune escape mechanism [34]. The attempt to develop about 100 anti-malaria vaccine candidates indicated their limited protective effect. The global analysis of antigens led to the conclusion that LCRs present in proteins containing glutamate-rich and/or repetitive motifs carried the most immunogenic epitopes. On the other hand the antibodies recognizing these epitopes appeared to be ineffective in an *in vitro* study [35]. In this context it seems to be very important to avoid the presence of LCRs in vaccine epitopes due to 1) low effectiveness of antibodies recognizing this region, even if LCRs are highly immunogenic and 2) the presence of LCRs in some human proteins.

## Results

Our aim was to find the LCRs in the SARS-CoV-2 genome that are similar to fragments of human proteins and to identify if any of those overlap with other epitopes in an attempt to eliminate epitope hits that are too similar to the human proteome fragments.

### Protein similarity between SARS-CoV-2 and human LCRs

To achieve this goal we used our three methods: GBSC, MotifLCR and LCR-BLAST. GBSC takes as an input whole protein sequences, however the input for LCR-BLAST and MotifLCR is expected to consist of LCRs. Therefore to identify LCRs in the SARS-CoV-2 and in the human proteomes we used the SEG tool with default parameters. There are 23 LCRs in SARS-CoV-2 proteome that were found in the following proteins: nsp2 (636 aa, one LCR found), nsp3 (1945 aa, six LCRs found) nsp4 (500 aa, one LCR found), nsp6 (290 aa, one LCR found), nsp7 (83 aa, two LCRs found), nsp8 (198 aa, two LCRs found), S protein (1273 aa, two LCRs found), E protein (75 aa, one LCR found), orf7a (121 aa, one LCR found), orf7b (43 aa, one LCR found), N protein (419 aa, four LCRs found) and orf14 (73 aa, one LCR found) (Fig 1). It is worth noting that most LCRs are located either in pp1a or in the C-terminal proteins (from spike to nucleocapsid protein). The middle section, from nsp9 to nsp16 is completely devoid of such sequences. In the next step we identified which of the SARS-CoV-2 LCRs are similar to human LCRs (Fig 1). Similar fragments are present in nsp3, spike glycoprotein and in the nucleocapsid protein. The list of these regions is presented in Table 1. We also provide a list of similar protein fragments from the human proteome obtained with three different methods (see S1-3 Tables).

**Figure 1.**
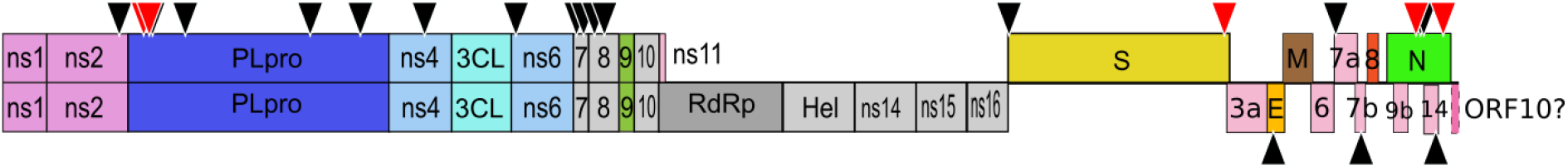
SARS-CoV-2 proteins shown according to their encoding position in the genome. Low Complexity Regions (LCRs) are marked with triangles. LCRs that are similar to human proteins are highlighted in red. The original figure for this modification was kindly provided by ViralZone, SIB Swiss Institute of Bioinformatics (https://viralzone.expasy.org/8996).

**Table 1.**
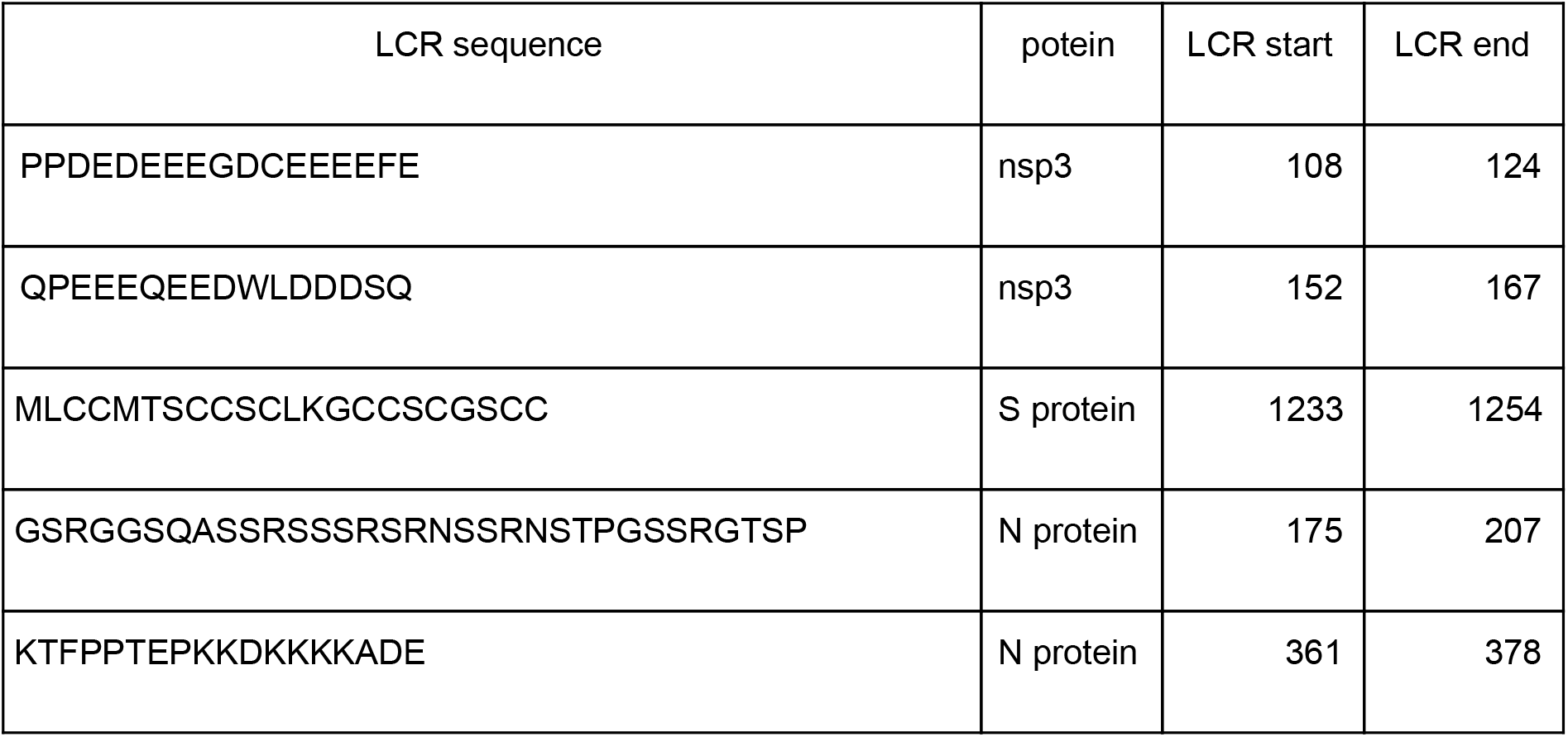
List of SARS-CoV-2 low complexity regions that are similar to human proteins.

Similarity of nsp3 is most significant to the myelin transcription factor 1-like protein (Myt1l) (Table 1, S1-S3 Tables). Myt1l was shown to be expressed in neural tissues in the developing mouse embryo [36]. Myt1l is supposed to limit non-neuronal genes expression, take part in neurogenesis and functional maintenance of mature neurons [37]. The glutamic acid-rich fragment is located close to the activation domain, however it was shown to be dispensable in this process [38].

The spike glycoprotein fragment MLCCMTSCCSCLKGCCSCGSCC has significant similarity to LCRs of ultrahigh sulfur keratin-associated proteins present both in hair cortex and cuticle (KRTAP 4.3, KRTAP 5.4 and KRTAP 5.9) [39] (Table 1, S2 and S3 Tables). KRTAPs are parts of the intermediate filaments of the hair shaft.

Nucleocapsid protein (N) has 2 LCRs that are similar to human LCRs (Table1, S1-S3 Tables) the most interesting comparable fragment is the zinc finger Ran-binding domain-containing protein 2 (RANB2), which is a part of the supraspliceosome where it is responsible for alternative splicing [40,41]. GBSC identifies the high similarity of the viral LCR to a LCR of the solute carrier family 12 which is an electroneutral potassium-chloride co-transporter which can be mutated in some severe peripheral neuropathies [42,43]. The C-terminal LCR is similar to a LCR from [F-actin]-monooxygenase MICAL3, actin-regulatory redox enzyme that directly binds and disassembles actin filaments (F-actin) [44]. This protein is also responsible for exocytic vesicles tethering and fusion, and cytokinesis [45–47]. The region of interest is probably involved in binding some of a multitude of binding partners of MICAL3 [46] (https://www.ebi.ac.uk/intact/interactors/id:Q7RTP6*).

Lists of human hits of LCRs similar to viral fragments were annotated with Gene Ontology (GO) terms [48] in order to find common functional features that were overrepresented among proteins composing the clusters. Here we focus on the results for GO annotations from the Biological Processes namespace since these functions may be crucial to understanding possible viral interventions into the cellular machinery. Complete lists of enriched GO terms are available in S4-S8 Tables. The best matches for the first LCR in nsp3 are human proteins involved in actin processing (S4 Table). The best matches for the adjacent LCR in nsp3 are related to signal transduction (S5 Table). The best hits for the spike protein LCR fragment are related to keratin (S6 Table). The human proteins similar to the central nucleocapsid protein’s LCR show discrepancies between sets of results. The output of GBSC clearly points to salinity response/salt stress responses (S7 Table). Results from LCR-BLAST and MotifLCR are actin-centred (S7 Table). In the case of the C-terminal nucleocapsid protein LCR, the most abundant human representatives are exocytosis and oxidation-reduction processes (S8 Table).

### Motif similarities of SARS-CoV-2 and human LCRs

We also tested similarities of viral LCR fragments to known domains and motifs using the UniProt, PROSITE, CDD, InterPro and ELM databases [49–53]. Most of the matches to known domains and motifs of the SARS-CoV-2 LCRs are to previously annotated regions, *i.e.* compositionally biased regions, rich in a particular amino acid or polyX regions. Only in two cases are there hits to specific domains.

The first similarity between SARS-CoV-2 LCR and a known motif is between the surface glycoprotein LCR (MLCCMTSCCSCLKGCCSCGSCC) and the keratin-associated protein domain (IPR002494). By using ScanProsite, we were able to find more than half a million of such motifs in the UniProtKB database [54]. Manual inspection of the viral LCR fragment shows the presence of a similar C-C-S-C motif. This fragment is also present in more than 500,000 sequences in UniProtKB. Interestingly, all 13 hits to the human proteome are metallothioneins with very similar motifs that are responsible for metal binding [55,56].

Nsp3 is the largest multi-domain protein encoded by the coronavirus genome. LCR of nsp3 (PPDEDEEEGDCEEEEFE) lies across the borders of two domains identified in coronaviruses: Ub1 (1-112) and acidic domain hypervariable region (HVR) (113-183)[57,58]. This LCR is significantly similar to the Armadillo-type fold (IPR016024), ‘a multi-helical fold comprised of two curved layers of alpha helices arranged in a regular right-handed superhelix, where the repeats that make up this structure are arranged about a common axis. These superhelical structures present an extensive solvent-accessible surface that is well suited to binding large substrates such as proteins and nucleic acids’ [59,60] https://www.ebi.ac.uk/interpro/entry/InterPro/IPR016024/.

### Non-recommended epitopes of SARS-CoV-2

In the last section we investigated the lists of epitopes suggested previously [22]; [16,61]. The authors of those papers provide predictions for 3295 possible candidates for T-cell epitopes and 1519 possible candidates for B-cell epitopes. By analysing this data we found that 21 of the predicted T-cell epitopes and 27 (1,7%) of the predicted B-cell epitopes overlap with 5 SARS-CoV-2 LCRs that are significantly similar to human proteins. However, only the S and N proteins from SARS-CoV are known to induce potent and long-lived immune responses [62–67]. This narrows the number of potential candidates to 562 (419 for S protein and 143 for N protein) for T-cell epitopes and to 397 (317 for S protein and 80 for N protein) for B-cell epitopes. Among these, we found out that 11 (2%) of the predicted T-cell epitopes and 19 (5%) of the predicted B-cell epitopes overlap with SARS-CoV-2 LCRs. The lists of B-cell and T-cell overlapping epitopes are presented in Tables 2 and 3 respectively and the overlapping fragments are marked in red colour. We therefore speculate that these regions should not be taken into account while selecting epitopes.

**Table 2.**
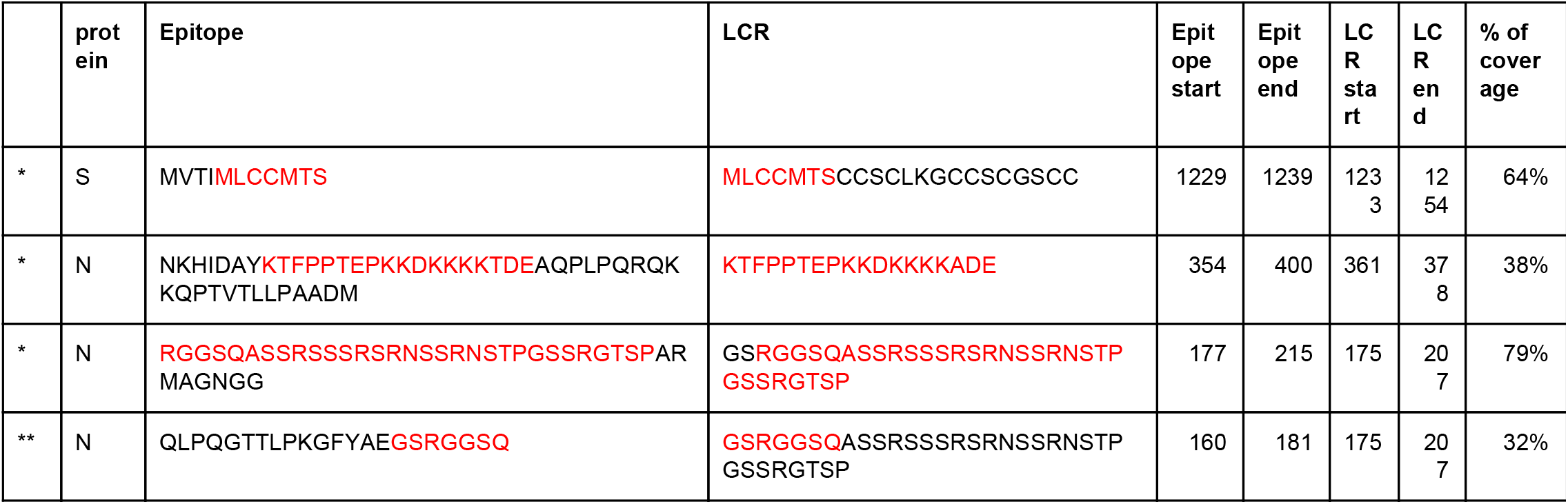
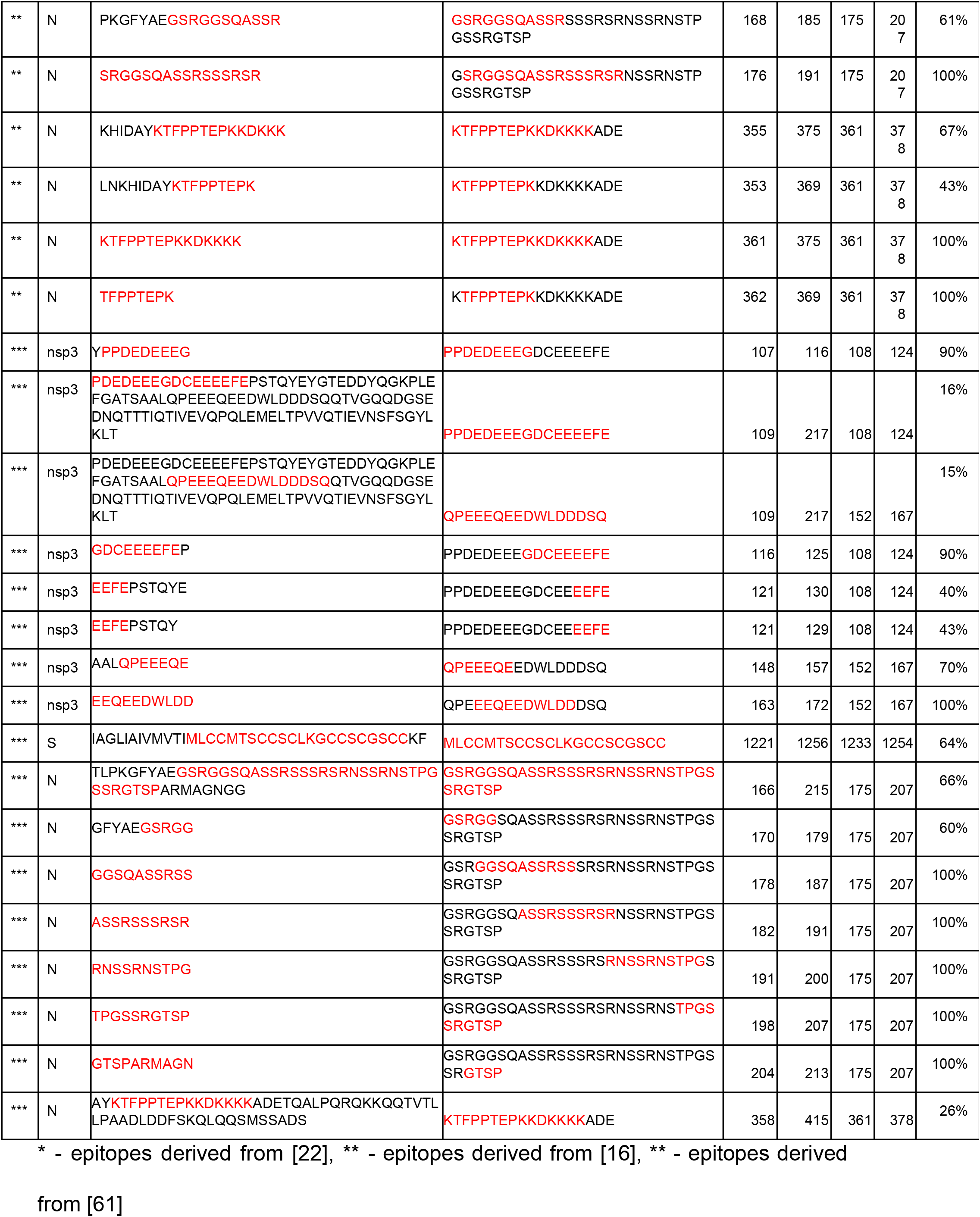
SARS-CoV-2 low complexity regions that overlap with B-cell epitopes.

**Table 3.**
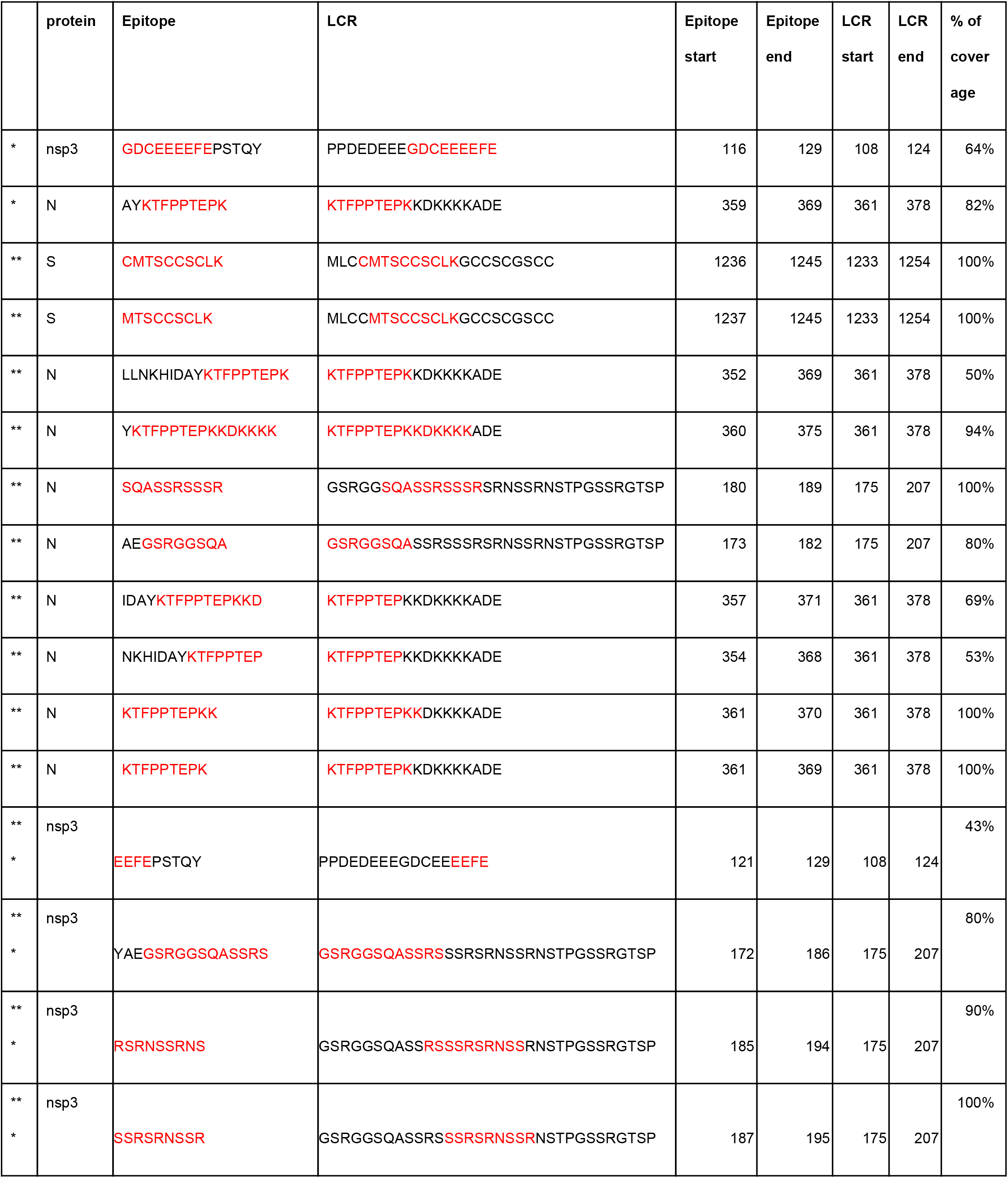
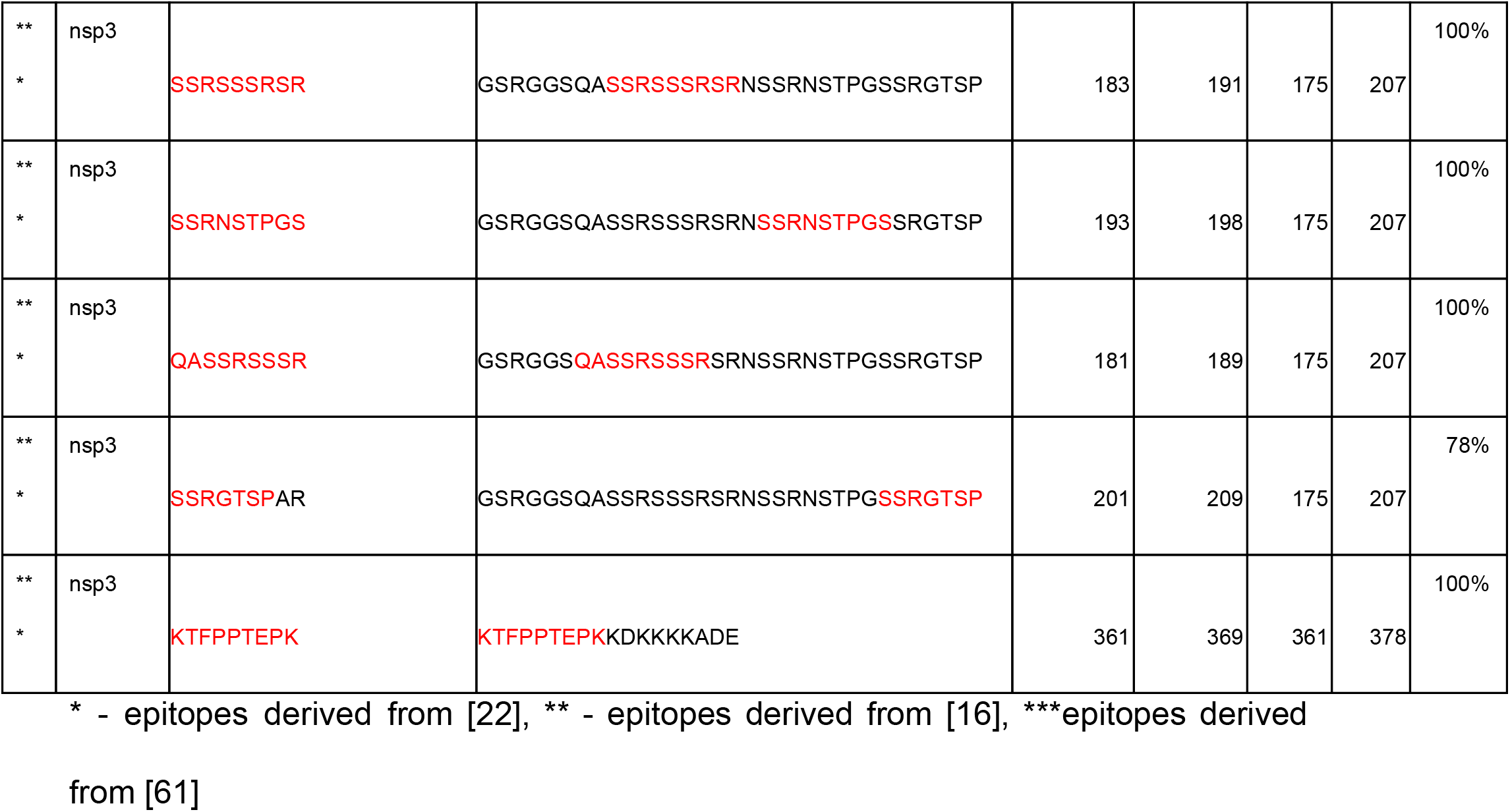
SARS-CoV-2 low complexity regions that overlap with T-cell epitopes.

## Discussion

Anti-COVID-19 vaccine development is mainly based on: DNA and RNA technology, peptides, virus-like particle, recombinant protein, viral vector, live attenuated virus and inactivated virus platforms [68]. Although, the epitopes for neutralizing SARS-CoV-2 antibody are known, the public information about the specific antigens which were used in vaccine development is not available. Some vaccines are based on S protein or even on whole virion [69]. Based on our findings in SARS-CoV-2 proteins 5 LCRs common for virus and human proteins are presented, therefore, it became obvious that antigens for SARS-CoV-2 vaccine development need to be designed and defined with extreme care. Based on vaccine development against SARS-CoV and MERS some concerns were recognized including induction of antigen-dependent enhancement (ADE) as not neutralizing antibody enhanced virus infectivity. ADE was found in cats vaccinated against a species-specific coronavirus [70]. In case of SARS, the use of whole inactivated virus or S glycoprotein induced hepatitis and lung immunopathology in animal models, while inactivated MERS in vaccination caused pulmonary infiltration in mice [71]. Moreover, it is still unclear whether adaptive T cell responses may also play a role in conferring protection against SARS-CoV-2. For SARS-CoV, in human survivors the memory T cells, but not B cells, were found around 6 years after infection [72]. The recent study indicated that in COVID-19 patient the 45 various antibodies against SARS-CoV-2 were found although only 3 exhibited ability to neutralize the virus [73]. Additionally, antibody against SARS-CoV-2 may cause the cross-reactivity with pulmonary surfactant proteins (shared similarity with 13 out of 24 pentapeptides) and development of SARS-CoV-2-associated lung disease [74]. Furthermore, last study indicated that antibody against S glycoprotein exhibited ability to cross-react with human tissue proteins including: S100B, transglutaminase 3 and 2 (tTG3, tTG2), myelin basic protein (MBP), nuclear antigen (NA), αmyosin, collagen, claudin 5+6 and thyroid peroxidase (TPO)[75].

Our work clearly shows similarity of SARS-CoV-2 protein low complexity sequences to human LCRs. We were able to detect similarity in 3 SARS-CoV-2 proteins to several human protein families. This resemblance can be seen in the nsp3, spike protein (S) as well as in the nucleocapsid protein (N). Previous research shows that both S and N proteins are known to induce potent and long-lived immune responses against SARS-CoV.

The nsp3 LCR fragments are part of the hypervariable region (HVR) which is Glu-rich. This region, even if so variable, is always present in all Coronoviridae. It is known to interact with nsp6, nsp8, nsp9 and its own C-terminal part, however no function has been assigned to it to date [58,76]. The same is true for the human transcription factor Myt1l’s glutamic acid-rich region which has an unknown role. Of note is the fact that the enrichment of glutamic acid was found as a feature of the highly immunogenic polypeptides [35]. Since Mytl1l is a transcription factor we may hypothesize that its LCR is somehow linked to the general function of binding nucleic acids. Such parallels may be helpful in understanding SARS-CoV-2 processes.

The surface glycoprotein (S) is of utmost interest to the scientific and medical communities because of its presence on the viral particle surface. The LCR identified in this study is a part of the cysteine-rich motif (CRM) present in the S2 domain, in the most C-terminal end of the protein located in the cytoplasm (endodomain) [77]. This sequence has been shown to be palmitoylated which is a critical step towards incorporation of S to the viral envelope [78–85]. Similarities to keratin-associated proteins and metallothioneins are hard to interpret. There are many possible explanations. One of them is the presence in epithelium. The function of this set of cysteines demands a more detailed study.

The nucleoprotein/nucleocapsid phosphoprotein (N) packages the viral genome into a helical ribonucleocapsid (RNP) and is crucial during viral self-assembly as shown in experiments with previously known coronaviruses [86,87]. Both regions of interest are located in the SR-rich region of the linkage region (LKR: residues 176–204) and the C-terminal disordered region (residues 370–389) that together with the N-terminal part are involved in RNA binding [88,89]. Similarity of the N protein to RANB2, an element of the supraspliceosome, seems surprising. However a hypothesis based on results from zebrafish may point at RANB2 as a weapon against infections, as is the case of the fish ZRANB2 [90]. The C-terminal LCR is similar to the human MICAL3 LCR which is multifunctional [45,46]. Gene Ontology analyses studies appear to indicate an intriguing over-representation of transport functions among human proteins whose LCRs are similar to coronavirus proteins.

It is known that viruses attack major cellular processes like vesicular trafficking, cell cycle, cellular transport, protein degradation and signal transduction to realize their goals [91]. Many host processes are taken over by viral proteins with the use of short linear motifs that are often parts of intrinsically disordered regions (IDRs). For example, the RGD motif mimics the regular cellular machinery for cell attachment *via* integrin [92]. Many IDRs are composed of low complexity regions. Therefore the hypothesis of the importance of similarities described above are not unfounded. Thorough analysis of SARS-CoV-2 short linear motifs has been recently published by the Gibson’s group [93].

The most important outcome of this work is the warning that epitopes cannot be selected based only on factors like phylogenetic conservation or potential epitope targets. For the safety of patients and procedures, all epitopes that may be similar to human proteome fragments should be discarded from further studies because the cure against SARS-CoV-2 may as well turn against the host.

Due to the fact that several research groups are working on the development of vaccine against SARS-CoV-2 it is very important to highlight its possible danger. The autoimmune diseases rate increased significantly in recent years. Moreover, it correlates with vaccination programmes [94]. Several studies indicated that vaccine components may induce autoimmune disease e.g. vaccine against Lyme disease can cause chronic arthritis and rheumatic heart disease [95]. However the mechanism triggering autoimmune disease after vaccination still remains unclear [96].

We also note a complete lack of LCRs in proteins originating from the pp1ab polyprotein (encoded by *orf1ab*), nsp9-nsp16 (Fig 1). Previous studies have shown that LCRs are more often present on protein ends [97], which are hard to define in polyproteins as in the case of pp1ab. The only distinguishing feature of these proteins is their function; most proteins from this group are involved in replication [98]. We speculate that the similarity of viral LCRs to human proteins may not be purely accidental but may be a molecular disguise. We suggest that SARS-CoV-2 may use these regions for specific functions that replace the cellular machinery for its own purposes.

Here we provide the scientific community with tools that allow the comparison of all types of low complexity fragments. These techniques have been shown to be useful previously in order to detect previously unknown similarities (Kubáň et al., 2019; Tørresen et al., 2019) and based on previous results we decided to use these tools to search for similarities among human and SARS-CoV-2 low complexity regions.

LCRs appear to come in 3 flavours. They can consist of homogenous polyX regions (homorepeats), repetitive fragments, or irregular LCRs [99]. Secondly, they usually come in specific combinations of amino acids, e.g. hydrophobic, cysteine-rich (alone or in combination with histidines), and glutamic acid always goes with aspartic acid. Our methods are tailored to detect the different types of such low complexity regions. The reader of this work should be aware that our results are based on sequence similarity only. We are fully aware that we do not include possible topological similarities of epitopes. These structural resemblances may of course play a role in comparison of even phylogenetically and fold-wise distant protein structures, as shown in allergic cross-reactivity [100,101]. We therefore cannot exclude that other similarities exist between SARS-CoV-2 and human proteins that are not identified here.

Overall, finding of five low complexity regions (LCRs) in three SARS-CoV-2 encoded proteins (nsp3, S and N) that are highly similar to regions from human proteome poses a serious threat to the vaccine or drug design. Similarity of SARS-CoV-2 LCRs to human proteins may have implications on the ability of the virus to counteract immune defense. The vaccine targeting LCRs may potentially be ineffective or alternatively lead to autoimmune diseases development.

## Materials and Methods

### SARS-CoV-2 protein sequences

All full-length protein sequences of the SARS-CoV-2 proteome were retrieved on 28 April 2020 from the ViralZone web portal (https://viralzone.expasy.org/8996) which provides pre-release access to the SARS Coronavirus 2 protein sequences in UniProt. The UniProtIDs of the SARS-CoV-2 proteins are P0DTC1 replicase polyprotein 1a (pp1a), P0DTD1 Replicase polyprotein 1ab (pp1ab), P0DTC2 Spike glycoprotein (S), P0DTC3 ORF3a protein (NS3a), P0DTC4 Envelope small membrane protein (E), P0DTC5 Membrane protein (M), P0DTC6 ORF6 protein, P0DTC7 ORF7a protein, P0DTD8 ORF7b protein, P0DTC8 ORF8 protein, P0DTC9 Nucleoprotein (N), P0DTD2 ORF9b protein, P0DTD3 ORF14 protein and A0A663DJA2 hypothetical ORF10 protein. Based on the information derived from UniProt replicase polyprotein 1a and replicase polyprotein 1ab were then divided into proteinases responsible for the cleavages of the polyproteins, that is: nsp1, nsp2, nsp3, nsp4, 3C-like proteinase, nsp6, nsp7, nsp8, nsp9, nsp10, nsp11, RNA-directed RNA polymerase, helicase, proofreading exoribonuclease and 2-O-methyltransferase.

### Identification of LCRs

To identify low complexity fragments in SARS-CoV-2 proteins we used the PlatoLoCo metaserver [31] which provides a web interface to a set of state-of-the-art methods that allow detection of LCRs, compositionally biased protein fragments, and short tandem repeats. Using all these methods we were only able to detect low complexity protein fragments using the SEG algorithm using the default parameters (*W* = 12, *K*1= 2.2, *K*2 = 2.5). To identify low complexity protein fragments in the human genome we downloaded human proteome from the Uniprot database (UP000005640) and analysed it using SEG with the same set of parameters.

### Searching for human protein fragments similar to SARS-CoV-2 LCRs

To detect human sequences that are similar to SARS-CoV-2 LCRs we used our three methods: GBSC, MotifLCR and LCR-BLAST (e-value threshold 0.001). The list of human LCRs that are similar to virus LCRs obtained with GBSC, MotifLCR and LCR-BLAST are presented in a S1, S2 and S3 Tables respectively.

For GBSC we used default parameters (score threshold 3, distance threshold 7). The method uses whole protein sequences as an input and then identifies repetitive regions that are consists of homopolymers or STRs. Then, similar protein fragments are clustered together and each cluster represents particular repetitive patterns. As a result we obtained two clusters that included both virus and human sequences.

MotifLCR and LCR-BLAST require low complexity fragments as input. In our case these sequences were obtained using the SEG tool as described above.

In the first step MotifLCR removes unique 2-mers in each sequence in order to create artificial sequences, then it searches for repeats in these new sequences and in the last step it creates clusters with native sequences that contain tandem repeats in artificial sequences. Repeat is defined as at least 3 times the occurrence of a specific amino acid pattern.

MotifLCR results consisted of 20 clusters that represented different repetitive motifs. However, the repetitive motifs in the obtained clusters were not specific. Therefore to further narrow down the sequences we used the results of MotifLCR as a subject database for LCR-BLAST and a list of viral LCRs as a query set. Finally, as a third tool we used LCR-BLAST with the viral LCRs as a query set and all human proteome LCRs as a subject database. As a result both MotifLCR and LCR-BLAST returned five clusters each with human LCRs sequences similar to SARS-CoV-2 LCRs.

### Comparing SARS-CoV-2 LCRs to epitopes

Having selected virus fragments that are similar to human sequences we then investigated the lists of T-cell and B-cell epitopes suggested by [22]; [16,61] in their works. The authors of the first work provide beginning and end amino acid coordinates for each epitope as well as a name of the virus protein and based on this information we were able to identify epitope regions that overlap with SARS-CoV-2 LCRs. In case of the list of epitopes provided by [16] and [61] we used WU-BLAST (http://blast.wustl.edu) with no gaps and parameters optimized for short sequences to find epitopes that align with 100% identity to SARS-CoV-2 LCRs and threshold of minimum length of aligned fragment of 4AA.

### Gene Ontology Enrichment

Gene Ontology enrichment functional analyses were performed on 12 clusters that included sequences similar to SARS-CoV-2 LCRs. Since some proteins may contain more than one LCR, and each of these LCRs may appear in the cluster, in order to avoid redundancy, enrichment analyses have been performed on lists of unique protein sequences. Reference sets for statistical analyses were created depending on the method used to generate clusters. In the case of GBSC to create a reference set we used all 11361 unique human proteins that composed all other clusters found by the method. In the case of MotfLCR as well as in the case of LCR-BLAST we used the same protein sets that were used to create bastp search databases and the sizes of the reference sets were 33880 and 45068 proteins respectively. To annotate human proteins with their corresponding GO terms from Biological Process, Molecular Function and Cellular Component namespaces we used BiomaRt R package [102]. Statistical analysis was performed with topGO R package [103] and to assess overrepresentation of GO term annotations in obtained clusters we applied hypergeometric test with false discovery Benjamin-Hochberg multiple testing correction with adjusted p-value cutoff 5%.

## Data and Code Availability

All data presented and analyzed in the present study was retrieved from the UniProt as described above. The published article includes all data and code generated or analyzed during this study, and they are summarized in the accompanying tables, figures and Supplemental Material.

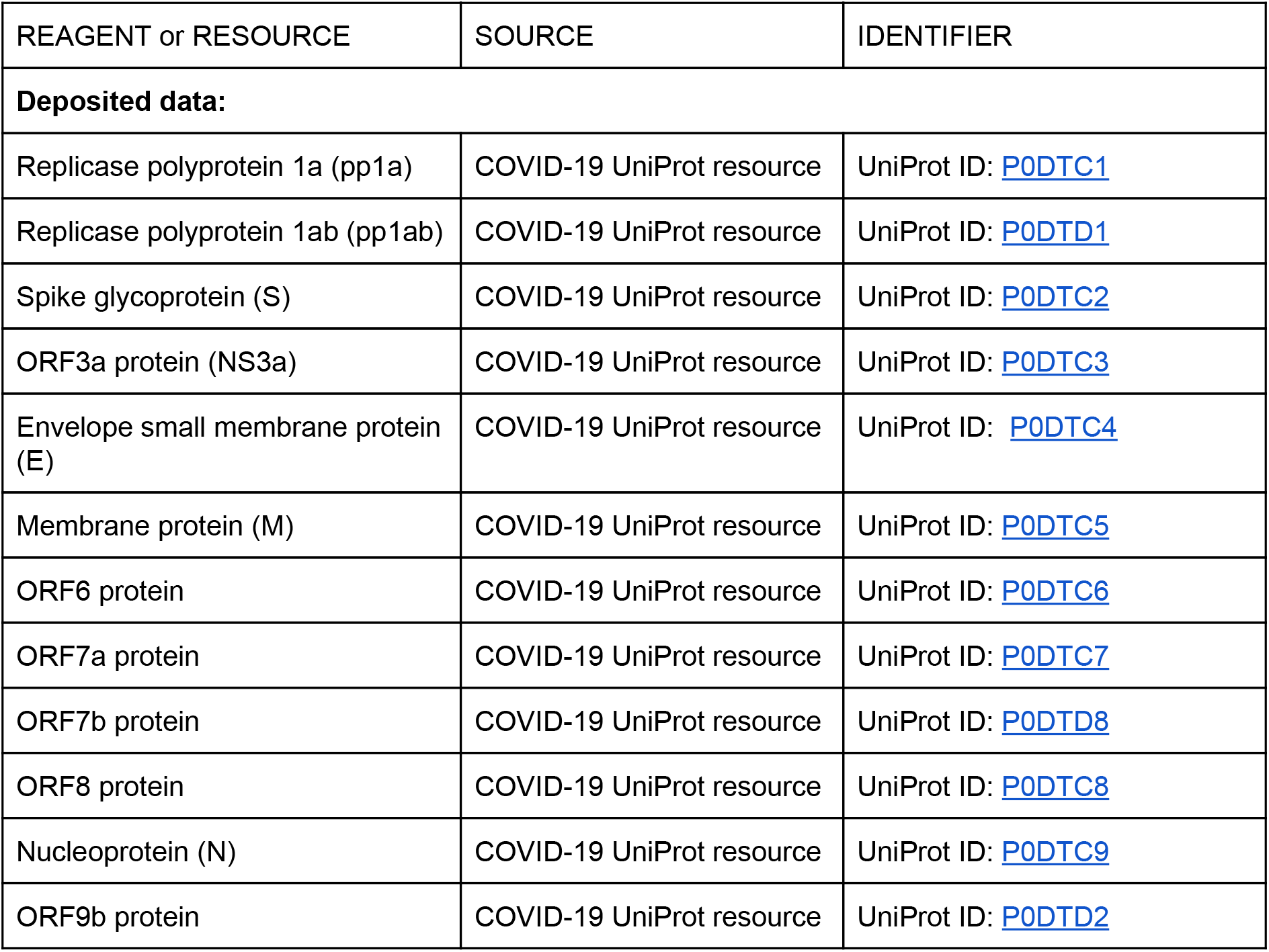
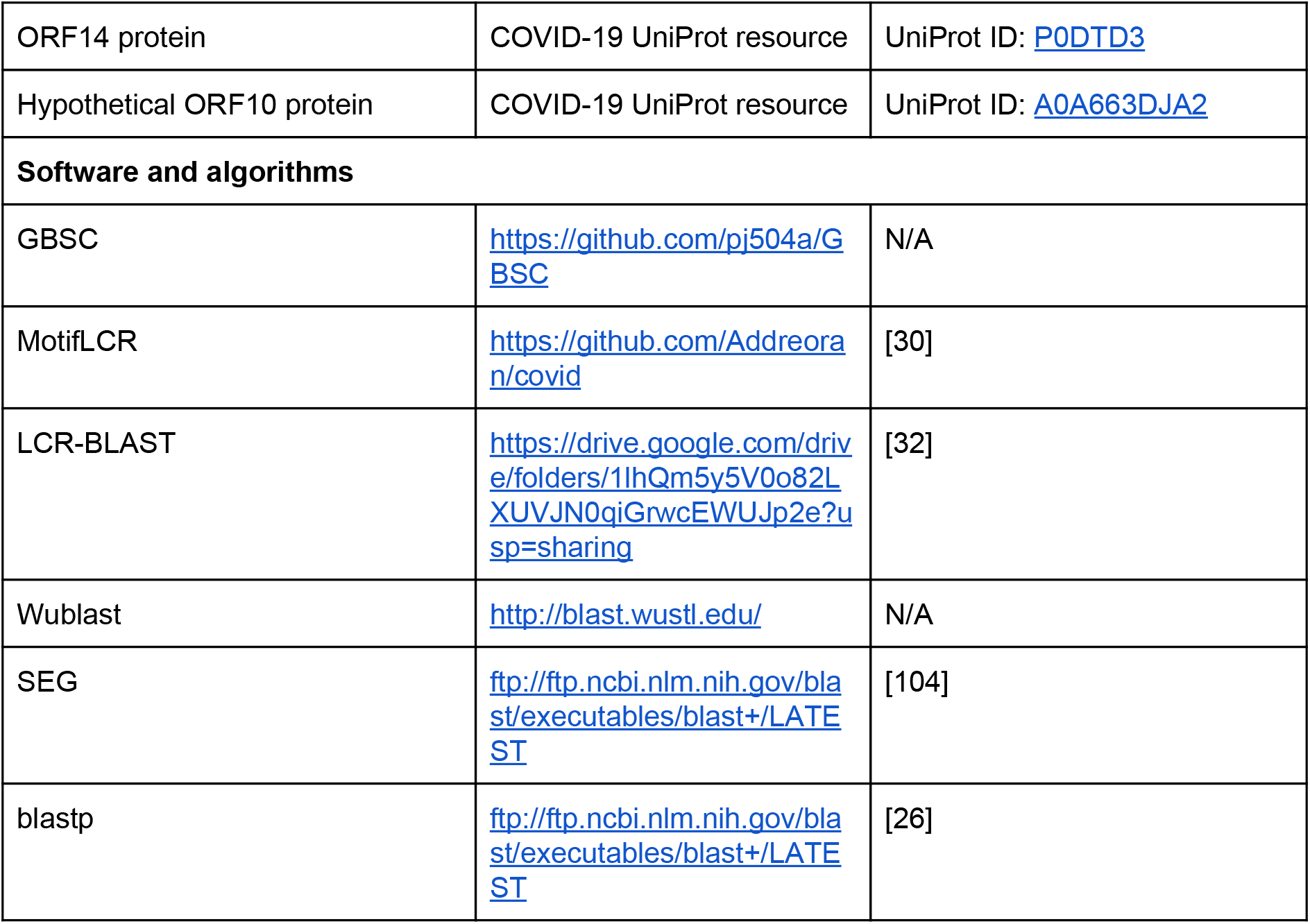

## Competing interests

We declare no conflict of interest.

## Acknowledgments

We thank Anna Muszewska, Eliana Kaminska, Miguel Andrade, Matthew Merski and Krzysztof Pawłowski for reading and commenting on the manuscript. We thank David Wotton for helpful discussions on Myt1l.

We are grateful to all the brave scientists and technicians for the acquisition of viral RNA and their analyses.

## Supplemental Information

**Supplementary Table S1**

S1 List of human LCRs similar to SARS-CoV-2 LCRs obtained from GBSC.

https://docs.google.com/spreadsheets/d/1JvGclCurlO6xTHgZNDl1o9HKbw6im4MHMGPcVm93buU/edit?usp=sharing

**Supplementary Table S2**

S2 List of human LCRs similar to SARS-CoV-2 LCRs obtained with MotifLCR -> LCR-BLAST.

https://docs.google.com/spreadsheets/d/1XhxTb11bbNdKWJq8OdUMUrF3-0-bZsogvU5bUWP92AE/edit?usp=sharing

**Supplementary Table S3**

S3 List of human LCRs similar to SARS-CoV-2 LCRs obtained with LCR-BLAST.

https://docs.google.com/spreadsheets/d/1xgifUaGyPe8uINe7gAaHsTqpHjiFSR_XdoHKPq-Nhik/edit?usp=sharing

**Supplementary Table S4**

S4 GO terms enriched for human proteins from all clusters obtained for the SARS-Cov-2 protein nsp3 LCR fragment 108-124.

https://docs.google.com/spreadsheets/d/1oGUIs2FC_6erG7-iuqdVIa9bMizNDlVLjawY8Eh5D48/edit?usp=sharing

**Supplementary Table S5**

S5 GO terms enriched for human proteins from all clusters obtained for the SARS-Cov-2 protein nsp3 LCR fragment 152-167.

https://docs.google.com/spreadsheets/d/1yxcPVAkKWoJTWHD4EREZDZWVOC_7g7IANtWaPP3J3NM/edit?usp=sharing

**Supplementary Table S6**

S6 GO terms enriched for human proteins from all clusters obtained for the SARS-Cov-2 spike glycoprotein (S) S LCR fragment 1233-1254.

https://docs.google.com/spreadsheets/d/19YVKIYOWcq2NGbNzvvKgdVHa71inO-w3ZLQStARp1XY/edit?usp=sharing

**Supplementary Table S7**

S7 GO terms enriched for human proteins from all clusters obtained for the SARS-Cov-2 nucleocapsid protein (N) LCR fragment 175-207.

https://docs.google.com/spreadsheets/d/1GkXBDXrNxTTyvpENecV4OrtIIOxgaSbI5xtYUuTBk7k/edit?usp=sharing

**Supplementary Table S8**

S8 GO terms enriched for human proteins from all clusters obtained for the SARS-Cov-2 protein N LCR fragment 361-378.

https://docs.google.com/spreadsheets/d/1rCuwIfHS9Ml9jwxTIS9-BOMkMVHeXAPG4V57wG048D0/edit?usp=sharing

